# Thoracic epidural blockade after myocardial infarction benefits from anti-arrhythmic pathways mediated in part by parasympathetic modulation

**DOI:** 10.1101/2024.03.14.585127

**Authors:** Jonathan D Hoang, Valerie YH van Weperen, Ki-Woon Kang, Neil R Jani, Mohammed A Swid, Christopher A Chan, Zulfiqar Ali Lokhandwala, Robert L Lux, Marmar Vaseghi

## Abstract

**Background:** Thoracic epidural anesthesia (TEA) has been shown to reduce the burden of ventricular tachyarrhythmias (VT) in small case-series of patients with refractory VT and cardiomyopathy. However, its electrophysiological and autonomic effects in diseased hearts remain unclear and its use after myocardial infarction (MI) is limited by concerns for potential RV dysfunction.

**Methods:** MI was created in Yorkshire pigs (*N*=22) by LAD occlusion. Six weeks post-MI, an epidural catheter was placed at the C7-T1 vertebral level for injection of 2% lidocaine. RV and LV hemodynamics were recorded using Millar pressure-conductance catheters, and ventricular activation-recovery intervals (ARIs), a surrogate of action potential durations, by a 56-electrode sock and 64-electrode basket catheter. Hemodynamics and ARIs, baroreflex sensitivity (BRS) and intrinsic cardiac neural activity, and ventricular effective refractory periods (ERP) and slope of restitution (S_max_) were assessed before and after TEA. VT/VF inducibility was assessed by programmed electrical stimulation.

**Results:** TEA reduced inducibility of VT/VF by 70%. TEA did not affect RV-systolic pressure or contractility, although LV-systolic pressure and contractility decreased modestly. Global and regional ventricular ARIs increased, including in scar and border zone regions post-TEA. TEA reduced ARI dispersion specifically in border zone regions. Ventricular ERPs prolonged significantly at critical sites of arrhythmogenesis, and S_max_ was reduced. Interestingly, TEA significantly improved cardiac vagal function, as measured by both BRS and intrinsic cardiac neural activity.

**Conclusion:** TEA does not compromise RV function in infarcted hearts. Its anti-arrhythmic mechanisms are mediated by increases in ventricular ERP and ARIs, decreases in S_max_, and reductions in border zone heterogeneity. TEA improves parasympathetic function, which may independently underlie some of its observed anti-arrhythmic mechanisms. This study provides novel insights into the anti-arrhythmic mechanisms of TEA, while highlighting its applicability to the clinical setting.

**Abstract Illustration:** Myocardial infarction is known to cause cardiac autonomic dysfunction characterized by sympathoexcitation coupled with reduced vagal tone. This pathological remodeling collectively predisposes to ventricular arrhythmia. Thoracic epidural anesthesia not only blocks central efferent sympathetic outflow, but by also blocking ascending projections of sympathetic afferents, relieving central inhibition of vagal function. These complementary autonomic effects of thoracic epidural anesthesia may thus restore autonomic balance, thereby improving ventricular electrical stability and suppressing arrhythmogenesis. DRG=dorsal root ganglion, SG=stellate ganglion.

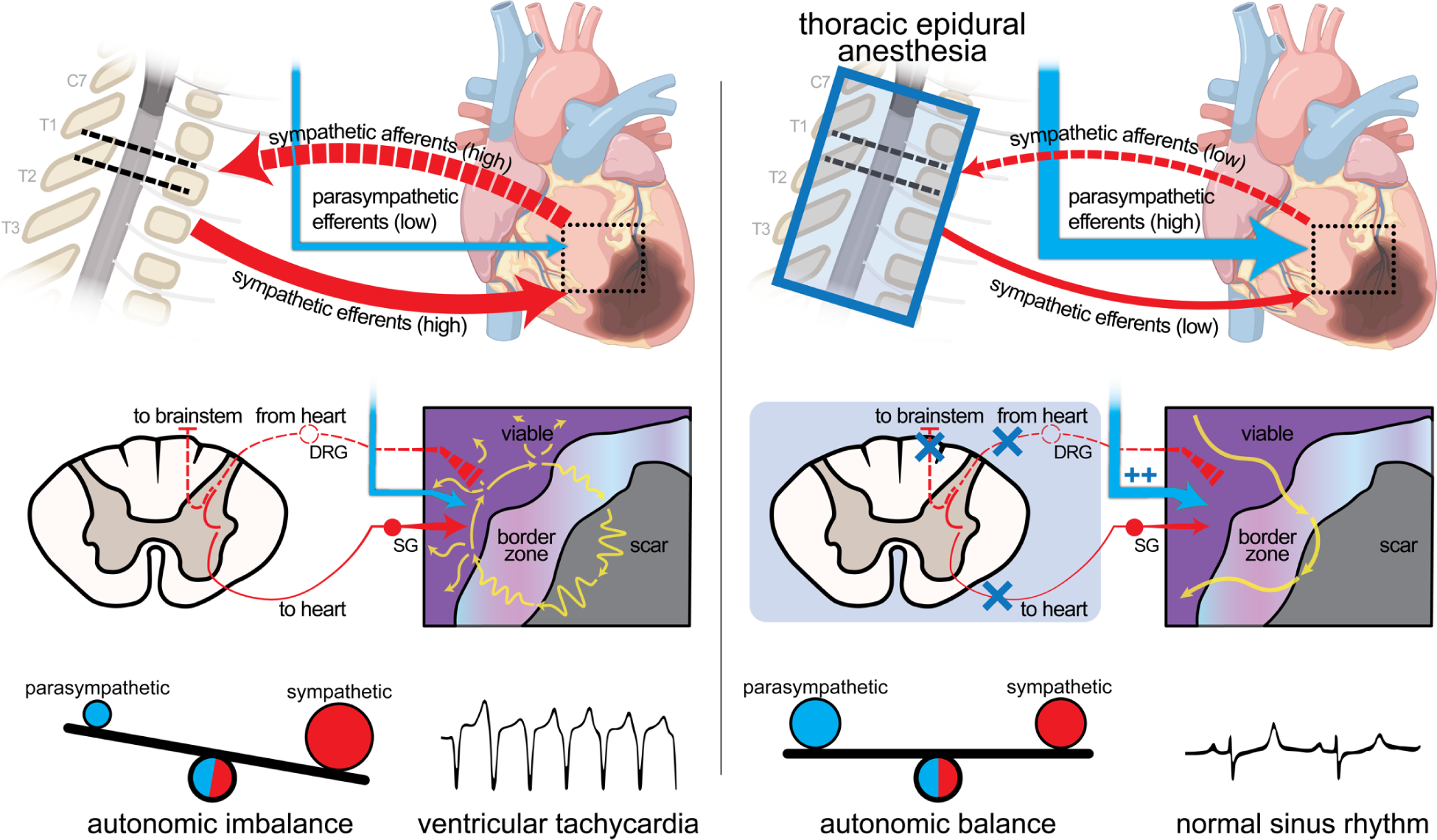

## INTRODUCTION

The sympathetic nervous system plays an important role in the occurrence of ventricular tachyarrhythmias (VT/VF).^1^ Cardiac sympathetic activation promotes triggered activity^2^ and shortens action potential duration and refractory periods,^3,4^ creating the trigger and substrate for VT/VF and leading to sudden cardiac death.^5^

Thoracic epidural anesthesia (TEA) acts at the dorsal and ventral horns of the spinal cord as a specific, but reversible, blockade of cardiac sympathetic efferent and afferent neurotransmission. Case reports and small case series of patients have shown that TEA may be effective in acutely decreasing the burden of refractory VT/VF in patients with structural heart disease and ventricular tachycardia (VT) storm.^6,7^ Previous experimental studies investigating the efficacy of TEA have been limited in scope and clinical relevancy by their application in normal hearts.^8,9^ In addition, the widespread adoption of TEA as an antiarrhythmic treatment has been limited by concerns and reports of possible right ventricular (RV) dysfunction, particularly in the setting of chronic left ventricular myocardial infarction (MI), where concerns for hemodynamic deterioration and preload dependence of the RV have limited the use of TEA in the setting of cardiomyopathy.^9^

To date, electrophysiological effects of TEA on the atria, ventricles, and the conduction system, its hemodynamics effects on right and left ventricular (LV) function, and its autonomic effects in diseased/infarcted hearts remain to be evaluated. Thus, the purpose of this study was to assess the biventricular hemodynamic, electrophysiological, and autonomic effects of TEA and evaluate underlying anti-arrhythmic mechanisms in the setting of chronic MI in a porcine model.

## METHODS

### Ethical approval

Twenty-two male Yorkshire pigs (S&S Farms) were used in this study. Care of animals conformed to the National Institutes of Health Guide for the Care and Use of Laboratory Animals. The protocol was approved by the University of California, Los Angeles, Institutional Animal Care and Use Committee.

### Study design

This was a prospective experimental in vivo porcine study with a repeated-measures design. Sample size calculations were based on a 50% assumed reduction in VT/VF inducibility (power>0.80, alpha=0.05) with TEA. Investigators were not blinded during the experimental analyses, however, all data was analyzed and corroborated by at least two independent researchers.

### Creation of myocardial infarction

MI was created percutaneously under fluoroscopic guidance as previously described.^10,11^ Briefly, animals (*N*=22; 39.6±0.6 kg) were sedated with tiletamine-zolazepam (4-8 mg/kg, intramuscular, IM), intubated, and general anesthesia maintained by isoflurane (1-2%, inhaled). Animals were treated intra-operatively with amiodarone (1.5 mg/kg, IM) and lidocaine (2 mg/kg, intravenous, IV) to reduce the incidence of iatrogenic ventricular arrhythmias. Next, a balloon-tipped coronary angioplasty catheter was advanced through a coronary guide catheter and over an angioplasty wire to the middle of the left anterior descending coronary artery (LAD) from the femoral artery. The balloon was inflated after the first diagonal branch of the LAD in each animal and 3 ml polystyrene microspheres (Polybead, 90 μm, Polysciences, Warrington, PA) were slowly injected via the lumen of the angioplasty catheter. The balloon was then deflated, and infarction was confirmed by lack of flow to the distal LAD on coronary angiogram coupled with ST-segment elevation. In the event of post-operative VT/VF, animals were defibrillated and administered an additional dose of amiodarone and lidocaine. Catheters and sheaths were then removed, the animal weaned from anesthesia, and monitored until ambulating.

### Animal preparation

Four to six weeks following MI, animals (53.6±1.4 kg) were sedated with tiletamine-zolazepam (4-8 mg/kg, IM) and general anesthesia maintained by isoflurane (1-2%, inhaled) throughout surgical preparation. Sheaths were placed in the bilateral femoral veins and arteries for saline infusion and pressure monitoring, respectively. Animals were transitioned to α-chloralose (50 mg/kg initial bolus, then 20-30 mg/kg/hr infusion, IV) for electrophysiological and autonomic assessment after the surgical portion of the study was completed.

### High thoracic epidural anesthesia

Pigs (*N*=16) were placed in the left lateral decubitus position. A 17-gauge Tuohy needle was inserted via the paramedian approach into the T5-T6 epidural interspace via standard loss-of-resistance approach and under fluoroscopic guidance (Figure 1A-C). A 19-gauge open-end epidural catheter (Teleflex Inc, Wayne, PA) was advanced beyond the needle tip into the epidural space. The catheter tip was advanced to the C7-T1 vertebral space and contrast injected to confirm appropriate placement under fluoroscopy. Lack of aspiration of blood and cerebrospinal fluid was used to exclude intravascular and intrathecal catheter placement. Lidocaine (2%, 0.2-0.4 ml/kg) was administered epidurally at a rate of 4 ml/min. Hemodynamic and electrical parameters were assessed immediately before (pre-TEA) and starting at 10 minutes after administration of lidocaine (post-TEA), to allow for stabilization of hemodynamic changes.

**Figure 1.**
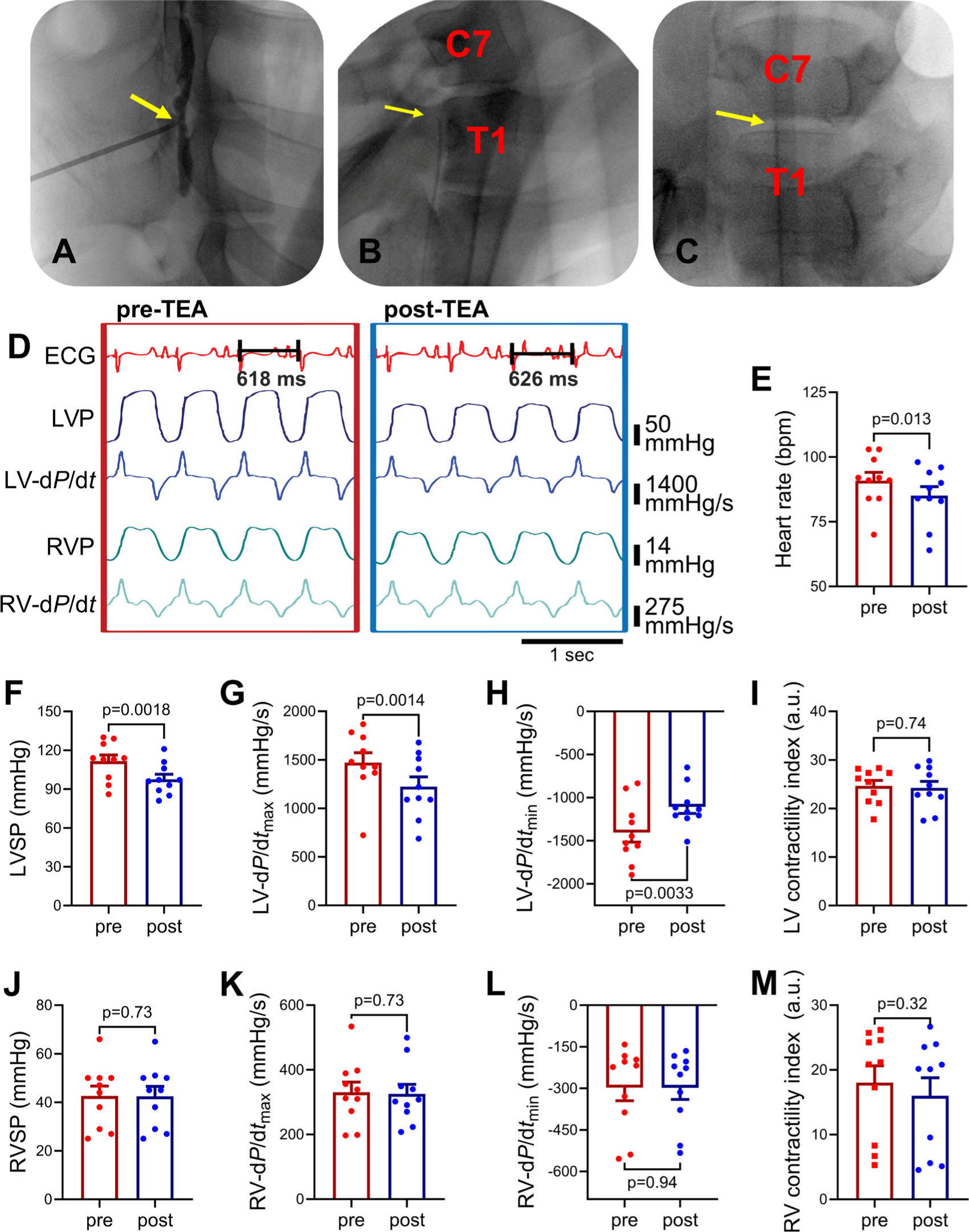
Effects of thoracic epidural anesthesia on hemodynamic parameters. **(A)** A 17-gauge Tuohy needle is advanced into the epidural space from the T5-T6 or T6-T7 interspace and contrast is injected to confirm the initial placement of the needle tip (arrow). **(B)** An open-tip epidural catheter (arrow) is placed with the animal in the lateral decubitus position and advanced from T5-T6 interspace to the C7-T1 vertebral level under fluoroscopic guidance. **(C)** Final catheter placement is confirmed in the supine position (antero-posterior view). **(D)** Representative raw traces of the effects of TEA on right ventricular (RV) and left ventricular (LV) function. Thoracic epidural anesthesia (TEA) modestly, but significantly, decreased **(E)** heart rate **(F)** LV systolic pressure (LVSP), **(G)** LV inotropy (LV-d*P*/d*t*_max_) and **(H)** LV lusitropy (LV-d*P*/d*t*_min_), but did not **(I)** decrease LV contractility index (LV-d*P/*d*t*_max_ • LVP^−1^). TEA did not have any effects on **(J)** RV systolic pressure (RVSP), **(K)** RV inotropy (RV-d*P*/d*t*_max_), **(L)** RV lusitropy (RV-d*P*/d*t*_min_), or **(M)** RV contractility index (RV-d*P/*d*t*_max_ • RVP^−1^). Pre-TEA *vs* post-TEA parameters were compared using Student’s paired *t*-test, *N*=10 animals.

### Cardiac electrophysiological recordings and analysis

Quadripolar pacing catheters were placed in the RV and right atrium (RA) to determine effective refractory periods (ERP) and measure atrio-His (AH) and His-ventricular (HV) conduction time (*N*=6). AH and HV intervals were measured pre- and post-TEA during atrial pacing at a drive cycle length (CL) of 450 msec.

Animals underwent median sternotomy to expose the heart. The pericardium was then opened and a pericardial sac created. A 56-electrode sock was placed over the ventricles in animals receiving TEA (*N*=15) to continuously record unipolar epicardial electrograms using a GE CardioLab System (Boston, MA). In a subset of animals (*N*=5), a constellation basket catheter (Boston Scientific, Marlborough, MA) was, under fluoroscopic guidance, advanced into the LV via the left carotid artery, allowing for the continuous recording of 64 local, endocardial, unipolar electrograms.

Activation recovery intervals (ARIs) were analyzed to estimate regional myocardial action potential durations,^12,13^ using customized software, iScaldyn (University of Utah, Salt Lake City, UT). ARIs were then mapped onto 2D-schematic polar maps with relative positions of anatomical landmarks noted to assess regional differences.

Bipolar epicardial voltage mapping was performed (*N*=15 animals) to delineate scar, border zone, and viable regions, using a standard electrophysiology catheter (2-2-2 duodecapolar catheter, Abbot, Minneapolis, MN). Briefly, the duodecapolar catheter was gently advanced between the epicardium and each sock electrode and the location of each electrode with respect to its underlying bipolar voltage was recorded. Using standard voltage criteria utilized in patients undergoing clinical electrophysiological procedures, regions were defined as either scar (0.05 mV<voltage<0.5 mV), border zone (0.5 mV<voltage<1.5 mV), or viable (voltage>1.5 mV).^14^ Of note, if an electrode was overlaying an area of very dense scar (voltage<0.05 mV) that did not give rise to a discernable or analyzable signal, data from this electrode was excluded from analysis.

### Ventricular hemodynamic measurements

A 5-Fr Millar pressure-conductance pressure catheter (Millar, SPR-350) was placed in the RV and LV (*N*=10) via the femoral vein and artery, respectively, for continuous pressure measurements throughout the experiment. Raw signals were digitized and recorded by CED Power1401 (Cambridge Electronic Design) and subsequently analyzed using Spike2. Instantaneous contractility was calculated for each animal from the first derivative of the ventricular pressure signal. Inotropy (d*P*/d*t*_max_) and lusitropy (d*P*/d*t*_min_) were calculated as the peak and trough of this curve, respectively. Inotropy was normalized to the instantaneous pressure at time of d*P*/d*t*_max_ to indirectly assess the load-independent effects of TEA on inotropy (contractility index; d*P*/d*t*_max_ • ventricular pressure^−1^).^15^

### Assessment of effective refractory period and slope of electrical restitution

Atrial and ventricular ERP were measured by extra-stimulus pacing (Micropace EP, EPS320) at a drive CL of 400-500 msec using an epicardial pacing catheter. An extra-stimulus was decremented by 5 msec down to ERP. The same drive CL and locations were chosen pre- and post-TEA. Ventricular ERPs were assessed at multiple sites: endocardially from the RV apex (*N*=10) and epicardially from the RVOT (*N*=10) and LV border zone (*N*=4; as defined by bipolar voltage mapping). Using the electrogram (EGM) closest to the origin of pacing, electrical restitution curves that described the relationship between ARI and preceding diastolic interval (DI) were constructed for epicardial pacing sites. ARI and DI were measured and fitted on a logarithmic curve. The maximum slope of a linear regression line fitted to the logarithmic curve, S_max_, was calculated for each ventricular site and compared pre- *vs* post-TEA.^16,17^

### Induction ventricular tachycardia and ventricular fibrillation

VT inducibility (*N*=10) was tested using programmed electrical stimulation, as is commonly performed in patients undergoing electrophysiological studies for VT (Supplemental Figure 1). Programmed stimulation was performed using an 8-beat drive train stimuli (S1) at delivered at a cycle length of 450 msec or 500 msec (depending on intrinsic sinus rate), followed by an S2 extra-stimulus. The same CL and pacing current were used pre- and post-TEA to minimize their effects on inducibility of VT. The extra-stimulus was set at ≥75% of S1 CL and decremented by 10 msec down to an S1-S2 interval of 200 msec or ERP, whichever came first. If the extra-stimulus reached ERP or the minimum coupling interval of 200 msec without causing sustained VT, 10-20 msec was added to the ERP interval to ensure ventricular capture, and the next extra-stimulus (S3) was added and then similarly decremented (by 10-20 msec until 200 msec or ERP). If sustained VT was not induced with S3, the S2-S3 coupling interval was prolonged by 10-20 msec to ensure stable capture of all preceding stimuli (S1-S3), before an S4 stimulus was added and decremented. Ventricular extra-stimulus testing was performed using Micropace (EP320) and Prucka CardioLab System (GE Healthcare) from the RV endocardium or LV anterior epicardium. VT/VF inducibility was defined as the occurrence of sustained VT (≥30 seconds) or VF requiring defibrillation. Sustained VTs were further classified as monomorphic or polymorphic VT based on VT morphology. Non-sustained VT (NSVT) was defined as ≥3 consecutive premature ventricular contractions, but lasting < 30 seconds. If VT was induced pre-TEA, the same ventricular site was used for VT inducibility post-TEA. Inducible animals were cardioverted if VT did not terminate after 30 seconds. If sustained VT was inducible at baseline, a minimum wait period of 60 minutes was allowed after cardioversion and before administration of TEA and repeat inducibility testing.

Susceptibility for arrhythmic events was also quantified via an arrhythmia score, taking into account both the number of extra-stimuli needed to evoke the arrhythmic event and the severity of the arrhythmic event.^18^ Briefly, 1 point was assigned for NSVT and 2 points for sustained VT (either monomorphic or polymorphic VT; MMVT or PMVT, respectively); this value was further scaled for the extra-stimuli at which the arrhythmias were observed (e.g., the ease of inducibility; S2=×3, S3=×2, S4=×1) and summed across all extrastimuli.^19,20^

### Evaluation of cardiac autonomic function

Bilateral stellate ganglia were isolated behind the parietal pleura and stimulated via bipolar needle electrodes (Natus).^10,21^ After bilateral lateral neck cutdowns, the cervical vagi were isolated and stimulated via bipolar spiral cuff electrodes (LivaNova, PLC). Threshold current was defined unilaterally for the stellate ganglia as the current causing a 10% increase in heart rate or systolic blood pressure at 4 Hz, 4 msec.^21,22^ Vagal thresholds were defined unilaterally at 10 Hz, 1 msec as the current causing a 10% decrease in heart rate.^10,21,23^ Bilateral stellate stimulation (BSS) was performed pre- and post-TEA at 4 Hz, 4 msec and 1.5× threshold current for 1 minute, to assess the effects on stellate-mediated sympathetic efferent activation (*N*=6). Right and left vagal nerve stimulation (VNS) were performed pre- and post-TEA at 10 Hz and pulse-width of 1 msec and 1.2× threshold current for 10 seconds to assess effects of TEA on VNS-mediated electrical and hemodynamic effects (*N*=6). A minimum of 30 minutes after BSS, and 10 minutes after VNS was allowed for recovery and stabilization of parameters.

Vagal baroreflex sensitivity (BRS) was tested by bolus intravenous administration of phenylephrine (3-5 μg/kg, intravenous) to evoke a 30-40 mmHg increase in systolic pressure. The same dose of phenylephrine was used before and after TEA (*N*=10). Sensitivity of the parasympathetic component of the baroreflex was determined as the slope describing the beat-to-beat relationship between increases in systolic blood pressure and RR interval, as described previously.^24^ Similarly, we administered the β-adrenergic receptor blocker metoprolol (5 mg, intravenous; average dose of 0.096±0.002 mg/kg) in a separate group of MI animals (*N=*6) to determine if the change seen in BRS with TEA were due to efferent sympathetic blockade (via blockade of β-adrenergic receptors). The same dose of phenylephrine (3-5 μg/kg) was used before and after β-adrenergic receptor blockade.

### Extracellular neural recording from the intrinsic cardiac nervous system

In-vivo extracellular neural recordings of the ventral interventricular ganglionated plexus (VIVGP), which is one of the intrinsic cardiac ganglia that innervates the ventricles, were obtained using custom-made 16-channel linear microelectrode arrays (MicroProbes for Life Science; 25 μm diameter platinum/iridium electrodes, 16 electrodes/probe, 375 μm interelectrode spacing; *N*=6). The probe was gently advanced into the epicardial fat pad at the atrioventricular junction, under the left atrial appendage, and connected to a 16-channel preamplifier (Model 3600, A-M systems, Sequim, WA, USA) with a head-stage pre-amplifier. Signals were sampled at 20 kHz, band-pass filtered (300 Hz to 3 kHz), digitized (Cambridge Electronic Design) and continuously recorded. Probe positioning was not changed pre- to post-TEA.

Neural signals were processed and analyzed offline using Spike2 software (Cambridge Electronic Design), as previously described.^25–28^ Artifacts were identified as simultaneous waveforms on all neural recording channels and subsequently removed. Individual neuronal spikes were identified using a threshold of 2× signal-to-noise ratio. Bulk activity of the VIVGP was measured as the sum of all neuronal firing of the 16 electrodes. Next, spike classification was performed using principal component, cluster on measurements, and K-means clustering analysis to identify unique neuronal waveforms. This allows for the neuronal firing pattern from a specific neuron to be followed over time and before/after interventions, given limitations of bulk firing analyses, which include firing from different types (afferent, efferent, convergent) neurons and/or ambient noise. Efferent post-ganglionic parasympathetic neurons in the VIVGP were identified based on their responses to VNS. Briefly, bipolar spiral cuff electrodes (LivaNova, PLC) were placed around each cervical vagi and the VIVGP activation threshold current, defined as the VNS current needed to evoke a 10% decrease in heart rate (20 Hz, 1 msec) was determined. Each cervical vagus was then stimulated separately for 1 minute at 1 Hz (1 msec, 1× VIVGP activation threshold current). This current and low frequency of stimulation have been shown to avoid hemodynamic changes that that can cause reflex neuronal responses, while still allowing for evaluation of changes in activity of intrinsic cardiac neurons to VNS.^25,28,29^ At least 20 minutes of recovery time was allowed between the two stimulations. Baseline activity during the 2 minutes before left or right VNS was compared to the two minutes upon start of stimulation (1 minute during VNS and 1 minute after) using the Skellam statistical test.^30^ Only neurons that showed significant changes in firing activity upon left or right VNS were identified as post-ganglionic parasympathetic neurons and included in the analyses. Basal activity (1 minute) was compared pre- and post-TEA.^25,28^ After identification of post-ganglionic parasympathetic neurons, neural activity of these specific neurons was followed and assessed for 20 minutes after infusion of lidocaine (post-TEA).

### Statistical analysis and reporting

Data are reported as mean±SEM. All analyses were performed as paired, two-tailed comparisons of pre- *vs* post-intervention (TEA or beta-blockade). No experiment-wide correction was applied. The Shapiro-Wilk test was used to confirm normality (if *N*≥6). Parametric testing was used for specific parameters with N<6, if prior studies suggested normal distribution of data (i.e., endocardial ARI and LV border zone ERP,^10^ and slope of restitution^31^). All remaining data for which normal distribution could not be confirmed were compared using non-parametric statistical testing, as detailed below.

Subsequently, paired two-tailed Student’s *t*-test was used to compare parameters pre-TEA *vs* post-TEA, including LV and RV hemodynamics, atrial and ventricular ERP, electrical restitution (S_max_), and AH and HV intervals. Global epicardial and endocardial ARIs were calculated as the average ARI of the 56 sock electrodes and 64 basket catheter electrodes, respectively. Regional epicardial ARIs were calculated as the average ARI of the electrodes classified by bipolar voltage mapping (4-5 electrodes per region) and compared pre-TEA *vs* post-TEA. The ARI of each electrode was subtracted from its post-TEA ARI, and these differences were averaged to compute the mean difference per region, and paired analysis of variance (ANOVA) was used to assess inter-regional differences in TEA-induced ARI changes. For responses to BSS and VNS, percent changes in global ARIs from baseline were calculated first; then paired two-tailed Student’s *t*-test was used to compare BSS- and VNS-induced ARI changes from baseline, before and after TEA. Maximal change in LVSP and HR were computed as the peak absolute change in each parameter induced by phenylephrine administration; these values were compared pre-TEA *vs* post-TEA and BL *vs* post-β-blockade by Student’s *t*-test. Bulk neural activity was compared pre-TEA *vs* post-TEA by the paired two-tailed Student’s *t*-test. Neurons significantly responding to VNS were identified by the Skellam test using R v4.3.2, as described above; the activity of VNS-responsive neurons were compared pre-TEA *vs* post-TEA by the paired two-tailed Student’s *t*-test.

Non-Gaussian parameters including arrhythmia score, pacing threshold, and electrical stimulation thresholds were compared pre-TEA *vs* post-TEA by the paired two-tailed Wilcoxon signed rank test. Transmural ARI gradient was assessed by subtracting the average ARI of the 64 basket catheter electrodes from the average ARI of the left ventricular epicardial sock electrodes (given the LV placement of the basket catheter). The transmural gradient was then compared by paired Wilcoxon rank signed test pre- *vs* post-TEA. Dispersion in AT/RT/ARI were calculated as the variance across global or regional electrodes. Baseline regional dispersion and regional change in dispersion were compared by paired Friedman test with post-hoc correction using the Benjamini Hochberg method. Change in dispersion (pre-TEA *vs* post-TEA) were compared by paired Wilcoxon signed rank test. BRS was computed as the slope defining the relationship between beat-by-beat RR-interval and LVSP; BRS was calculated for each individual test by simple linear regression,^24^ and linear regression resulting in non-significant correlations (P>0.05) and/or poor goodness of fit (r^2^<0.80) were discarded and tests repeated. BRS values pre-TEA *vs* post-TEA and pre-β-blockade *vs* post-β-blockade were compared by paired two-tailed Wilcoxon signed rank test. The exact binomial test (two-tailed) was used to compare VT inducibility pre-TEA *vs* post-TEA. P value<0.05 was considered statistically significant. All statistical analyses, unless otherwise stated, were performed with GraphPad Prism software v9.

All representative figures and images were selected from data points which visibly illustrated the methodology and qualitative observations while approximating the mean effect.

## RESULTS

### Hemodynamic changes resulting from high thoracic epidural blockade

To assess the effects of TEA on biventricular function, RV and LV function were continuously measured in infarcted animals before and after infusion of TEA (*N*=10, Figure 1A-D). Heart rate, LV systolic pressure, inotropy and peak rate of relaxation modestly decreased after TEA before stabilizing (HR: 90.9±3.2 to 85.1±3.5 bpm, p=0.013, LVSP: 111.6±4.7 to 97.6±3.8 mmHg, p=0.0018; LV-d*P*/d*t*_max_: 1472±102 to 1224±100 mmHg/s, p=0.0014; LV-d*P*/d*t*_min_: −1405±112 to −1107±76 mmHg/s, p=0.0033), Figure 1E-H. The LV contractility index (d*P*/d*t*_max_ normalized for systolic pressure; LV-d*P/*d*t*_max_ • LVP^−1^) showed no significant differences pre- *vs* post-TEA, suggesting that these changes in d*P*/d*t* are primarily driven by decreases in aortic systolic pressure, and not diminished LV contractility, Figure 1I. However, despite changes in LV hemodynamics, no effect on RV systolic pressure (42.6±4.1 to 42.5±4.0 mmHg; p=0.73) or RV inotropy (331±32 to 325±30 mmHg/s; p=0.73) was noted, Figure 1J-K, and RV lusitropy also remained unchanged, Figure 1L. Similar to the LV, the RV contractility index (RV-d*P/*d*t*_max_ • RVP^−1^) was also unchanged, Figure 1M. Importantly, RV function was unaffected despite its preload dependence, potentially through a balanced reduction in heart rate and establishment of more favorable right ventricular filling time. Hemodynamic instability or deterioration was not observed.

### TEA mitigates inducibility of ventricular arrhythmias

VT inducibility was assessed by ventricular extra-stimulus pacing before and after TEA (*N*=10), Figure 2A-B. All ten infarcted animals tested (100%) were inducible for sustained MMVT or polymorphic VT/VF requiring defibrillation pre-TEA. After TEA, inducibility was assessed at the same site from which VT/VF had been induced pre-TEA. Only 3 of 10 (30%) infarcted animals remained inducible for sustained VT/VF post-TEA. We further analyzed the incidence of non-sustained VT (NSVT), monomorphic VT (MMVT), and polymorphic VT (PMVT), Figure 2C-F. TEA decreased overall inducibility of VT/VF by 70% (p<0.0001, Figure 2G); the incidence of NSVT and sustained MMVT was also significantly decreased post-TEA. This was reflected in a significant decrease in the arrhythmia scores pre- *vs* post-TEA (pre-TEA 3.7±0.5 *vs* post-TEA 0.7±0.3, p=0.0078), Figure 2H.

**Figure 2.**
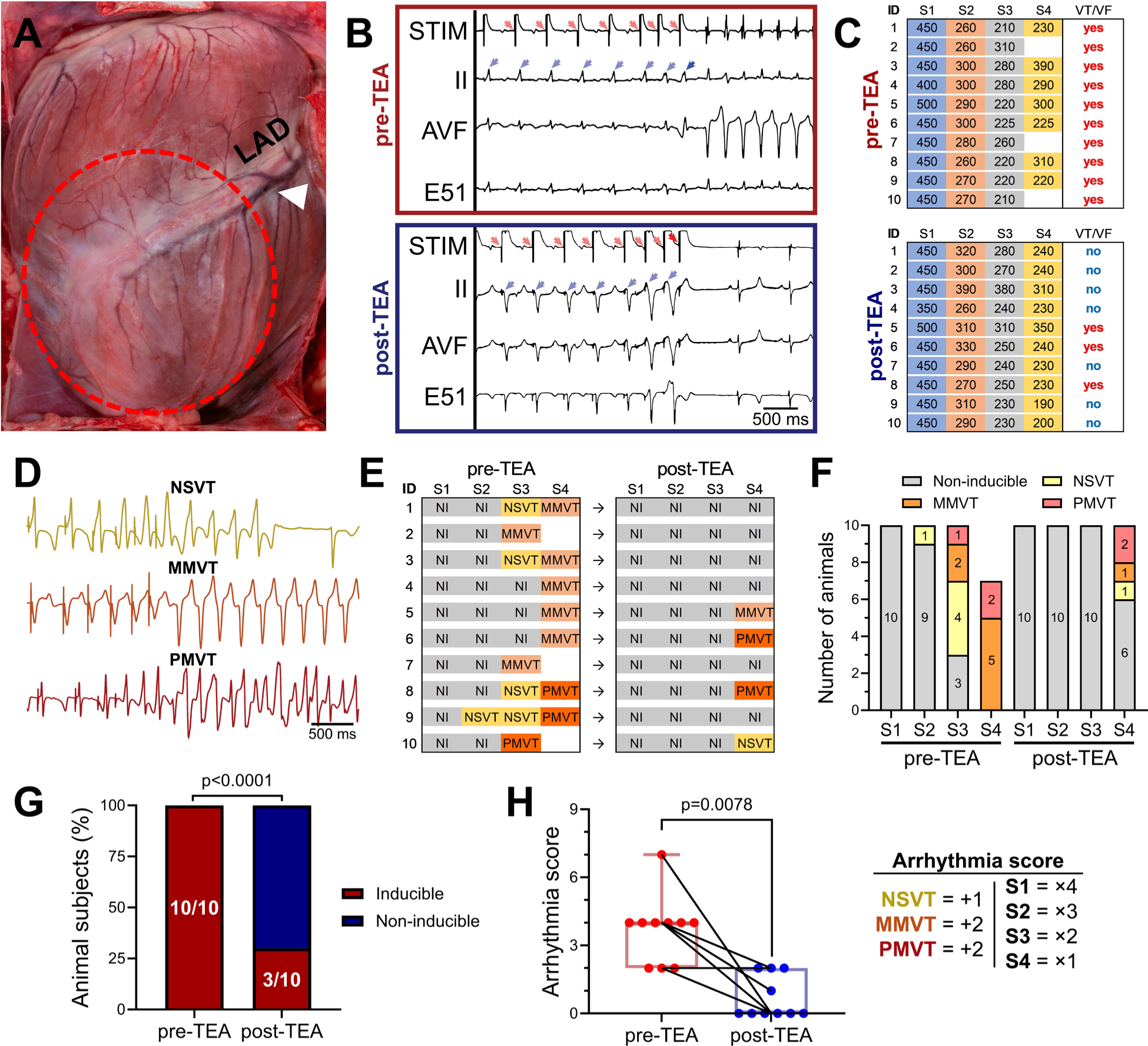
Effects of thoracic epidural anesthesia on electrical stability and inducibility of ventricular tachyarrhythmias. **(A)** Myocardial infarcts were created by occlusion of the left anterior descending coronary artery (LAD) immediately after the first diagonal branch (white arrowhead). The infarcted ventricular region, six weeks post-MI, is indicated by the red dashed line. **(B)** Example of VT/VF induction using programmed ventricular stimulation. Prior to thoracic epidural anesthesia (TEA), this animal was inducible for ventricular tachycardia (VT) with double extra-stimuli. After TEA, VT was no longer inducible with triple extra-stimuli, and ventricular effective refractory period (ERP) was reached (red arrows indicate pacing artifact and blue arrows indicate ventricular capture). **(C)** Breakdown of final stimulation parameters used to induce VT. **(D)** Examples of ventricular arrhythmias induced by programmed ventricular stimulation and **(E-F)** the frequency of their occurrence with each extra-stimulus. **(G)** Before TEA, all ten animals were inducible for VT, including 3 animals that were inducible with only 2 extra stimuli. After TEA, only 3 animals were inducible with up to 3 extra-stimuli, overall decreasing VT/VF inducibility by 70%. **(H)** An arrhythmia score summarizing the severity of the arrhythmia and the ease of inducibility further highlights the increase in electrical stability after-TEA. E51=cardiac electrograms from sock electrode #51; STIM=channel used for ventricular stimulation pacing. VT inducibility pre-TEA *vs* post-TEA was compared by the paired exact binomial test. Arrhythmia score pre-TEA *vs* post-TEA compared by the paired Wilcoxon Rank Sum test; *N*=10 animals.

### Supraventricular electrophysiological effects of TEA

Given mixed results of clinical studies on TEA’s effects on post-operative atrial fibrillation incidence,^32,33^ we first evaluated the electrophysiological effects of TEA on both the atrium and the conduction system in our model. Atrioventricular conduction time was assessed by measuring the atrial to His bundle conduction time (AH interval; *N*=6). TEA significantly prolonged the AH interval (92±6 to 108±5 msec; p<0.0035), without affecting the HV interval, despite atrial pacing at the same cycle length pre- and post-TEA (Supplemental Figure 2). Next, the effects of TEA on atrial refractoriness were assessed by extra-stimulus pacing of the left atrium (*N*=10). Importantly, TEA significantly prolonged atrial ERP (147±5 to 168±5 msec; p<0.0001), suggesting a potential benefit for supraventricular arrhythmias in the setting of MI-induced autonomic remodeling, Supplemental Figure 2.

### Effects of TEA on myocardial action potential duration and refractoriness

We next sought to determine the ventricular electrophysiological effects of TEA in chronically infarcted animals (*N*=15), including on specific regions, as classified by voltage mapping, Figure 3A. TEA induced a gradual prolongation of ventricular ARIs that stabilized within the first 90 seconds of infusion, Figure 3B. While TEA did not alter ventricular conduction (Supplemental Figure 3), TEA significantly prolonged global ventricular ARIs by 14±5 msec (354±8 to 375±7 msec; p=0.0008), Figure 3C-E. Importantly, bipolar voltage mapping to delineate regions of viable myocardium from scar and border zone regions revealed that this effect was not restricted to just the viable myocardium, Figure 3F. While ARIs from viable regions of myocardium were prolonged by 25±6 msec (358±9 to 383±8 msec; p=0.0016), ARIs of both scar (356±11 to 374±10 msec; p=0.0042) and border zones (361±8 to 387±9 msec; p=0.0047) also significantly prolonged, Figure 3F. There was no difference in the mean prolongation of regional ARIs, Figure 3G. Prior to TEA, border zone regions demonstrated the greatest dispersion in ARI, Figure 3H. While TEA had no effect on dispersion of ARIs within scar and viable regions, it significantly reduced ARI dispersion in the border zone region by nearly 50% (from 537±51 ms^2^ pre-TEA to 310±82 ms^2^ post-TEA; p = 0.025), Figure 3H-I.

**Figure 3.**
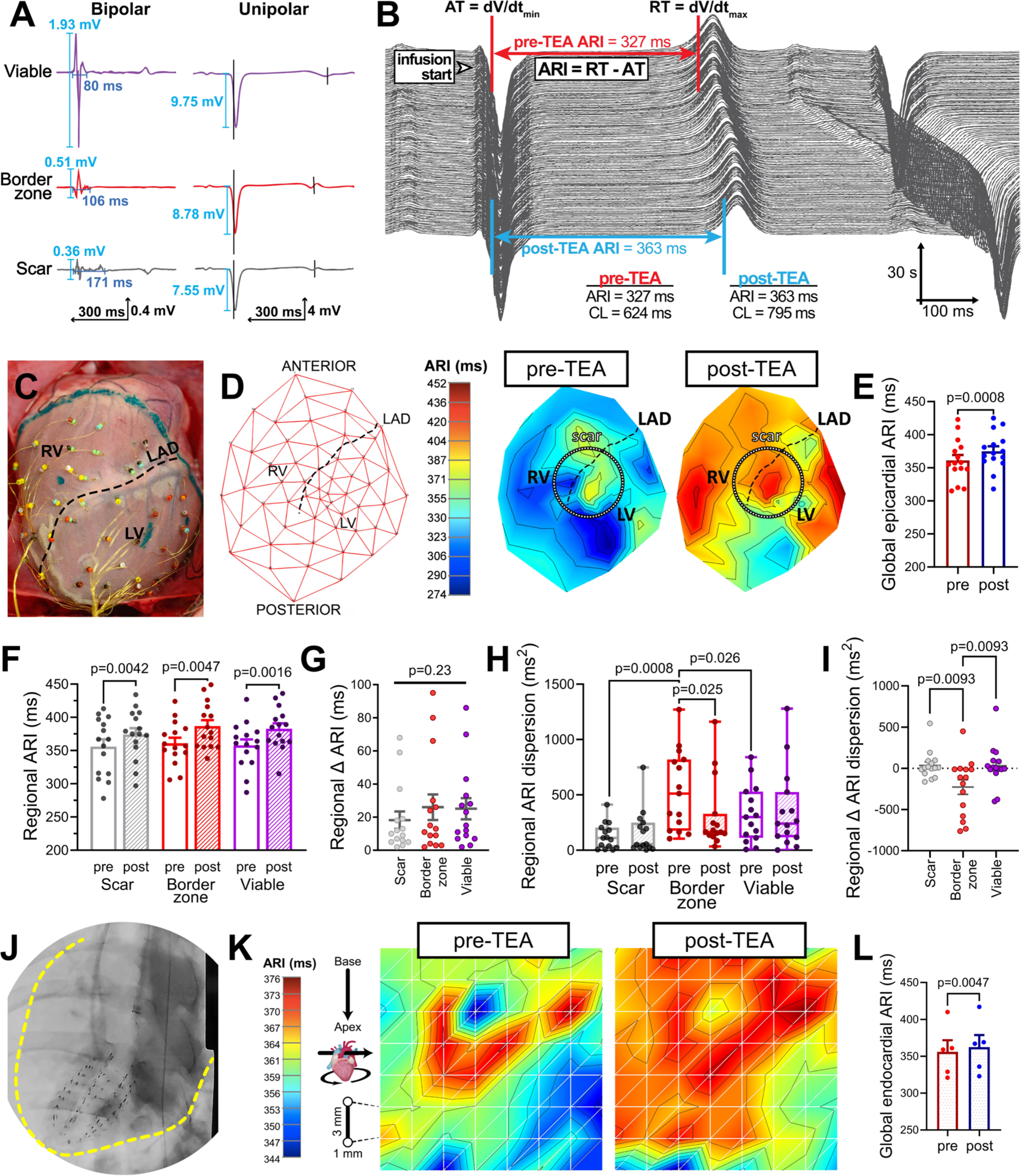
Effects of thoracic epidural anesthesia on ventricular action potential duration. **(A)** Representative epicardial bipolar electrograms (EGMs) and their corresponding unipolar EGMs from scar, border zone, and viable myocardium. **(B)** Representative waterfall plot of a unipolar electrogram from a single electrode in one animal spanning the time frame from a few seconds of baseline (before lidocaine administration) followed by the 90 second period after of lidocaine infusion. Over this period, there is progressive prolongation of activation recovery interval (ARI), which then plateaus. ARI, a surrogate for local action potential duration, is calculated as the time from activation time (AT; d*V*/d*t*_min_ of activation wave-front) to recovery time (RT; d*V*/d*t*_max_ of repolarization wave-front) from each unipolar electrogram. **(C)** A 56-electrode sock is placed around the ventricles to record local unipolar EGMs. **(D)** Electrograms were mapped onto a 2-D polar map for the assessment of global and regional electrophysiological changes. Representative epicardial polar maps comparing ARIs pre-TEA and post-TEA. **(E-G)** Global and regional ARIs from scar, border zone, and viable areas significantly prolonged post-TEA with similar prolongation seen between regions. **(H)** Regional ARI dispersion was greatest in border zone regions, reflective of greater heterogeneities in action potential duration in this region compared to scar and viable regions. TEA decreases the ARI dispersion in border zone regions **(I)** The reduction in border zone dispersion was significantly greater than either scar or viable regions. **(J)** A 64-electrode Constellation catheter was advanced under fluoroscopic guidance into the LV to measure local endocardial unipolar electrograms. The dashed yellow line marks the border of the ventricular wall. **(K)** Mapping the endocardial electrograms onto a 2-D polar map shows the global prolongation of endocardial ARI in the LV. **(L)** Endocardial ARIs significantly prolonged post-TEA. Pre-TEA *vs* post-TEA global, regional, and LV-specific ARIs were compared by Student’s paired *t*-test. Inter-region changes in ARI were compared by repeated-measures ANOVA. Baseline regional dispersion and regional change in dispersion were compared by paired Friedman test with post-hoc correction (non-significant comparisons not shown). Change in dispersion (pre-TEA *vs* post-TEA) were compared by paired Wilcoxon signed rank test. *N*=15 animals for global and regional epicardial data. *N*=5 for endocardial data.

To next investigate if these epicardial changes were conserved at the endocardium, we measured local endocardial EGMs from the left ventricle (*N*=5), Figure 3J-K. Like the epicardium, endocardial ARIs also significantly prolonged (356±16 msec to 362±16 msec, p=0.0047), Figure 3L. TEA did not significantly alter LV global epicardial or endocardial dispersion of ARIs, Supplemental Figure 3. With simultaneous epicardial sock and endocardial basket recordings (*N*=5) we further assessed the effects of TEA on transmural ARI gradients, but found no significant effect on these gradients pre- *vs* post-TEA, Supplemental Figure 3.

To determine TEA-induced modulation of tissue excitability, we assessed ventricular effective refractory period (ERP) at multiple sites by programmed electrical stimulation, Figure 4A-B. Pacing thresholds, defined as the current needed to evoke a paced beat, were unaltered by TEA, Figure 4C; however, TEA significantly prolonged ventricular ERP (262±8 to 292±10 msec; p=0.0057) as assessed endocardially from the RV apex (Figure 4D), a standard site for VT inducibility testing in the clinical setting.^34^ ERP was also increased in two other tested sites, including the right ventricular outflow tract (RVOT, Figure 4E), and in the left ventricular border zone (Figure 4F), regions shown to be critical to the genesis in ventricular arrhythmias.^35–37^ Electrical restitution curves were constructed from EGM recordings of the RVOT and LV border zone, Figure 4G-H. S_max_, the maximal slope of the curve describing the relationship between ARI and the preceding DI, was significantly decreased post-TEA; RVOT S_max_ decreased from 1.3±0.2 to 0.3±0.3 a.u. (p=0.0026), and LV border zone S_max_ decreased from 2.0±0.4 to 0.7±0.1 a.u. (p=0.046), Figure 4I-J.

**Figure 4.**
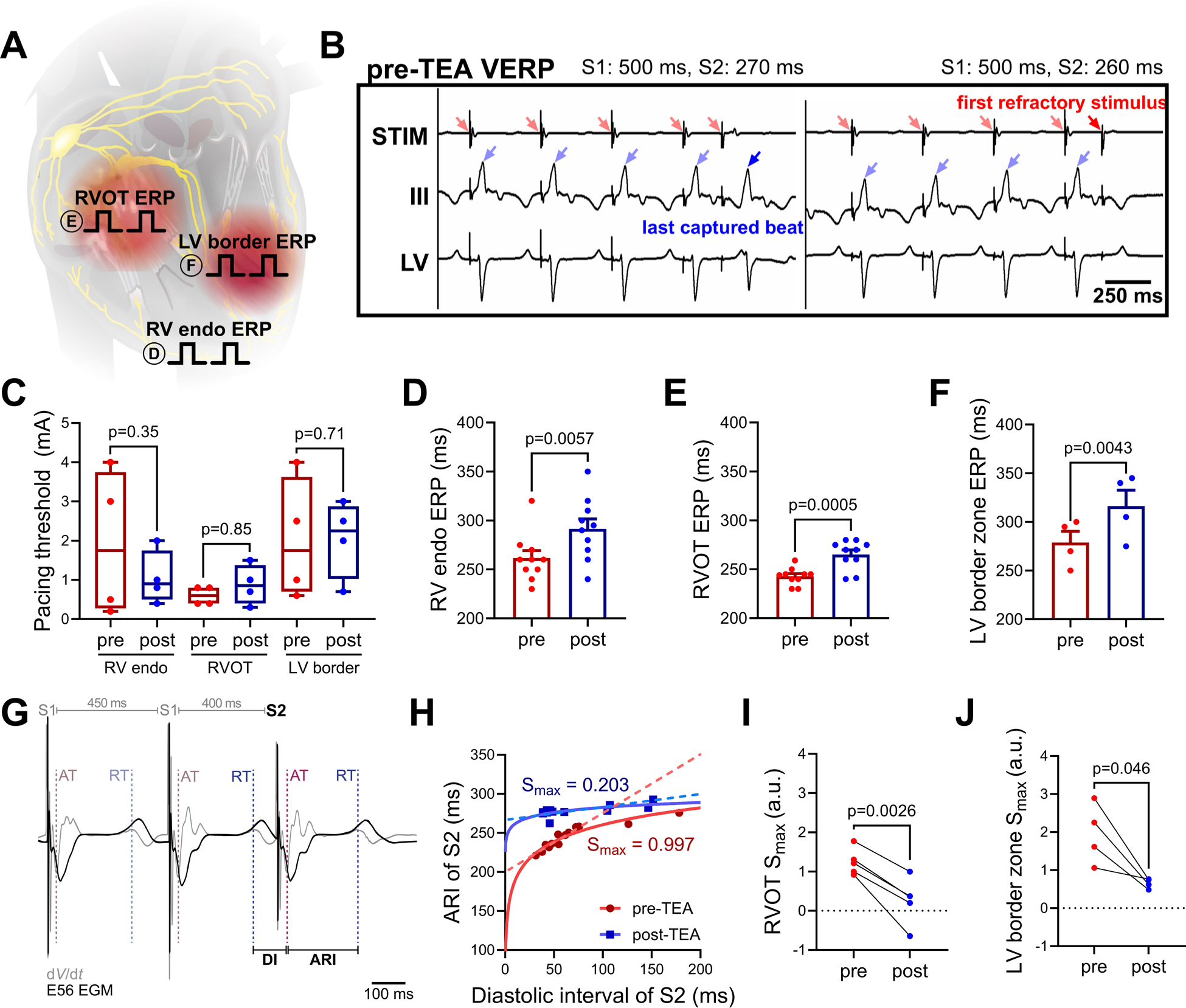
Effects of thoracic epidural anesthesia on ventricular refractory periods and restitution. **(A)** Graphical representation of pacing sites for measurements of effective refractory period (ERP) at the right ventricular (RV) endocardium, RV outflow tract (RVOT, epicardium) and epicardial left ventricular border zone region (LV border). **(B)** Example of ventricular ERP (VERP) measurements from one animal from the RVOT (pre-TEA), illustrating the last captured beat and the loss of capture after a further reduction in the coupling interval of S2. **(C)** TEA did not affect the pacing thresholds for ventricular ERP testing, but did prolong VERP at the **(D)** RV endocardium, **(E)** RVOT, and **(F)** LV border zone. **(G)** Local minima and maxima of the d*V*/d*t* of the EGM corresponding to site of pacing were assessed to measure the AT and RT of each beat, respectively. The diastolic interval (DI) was calculated as the difference in the RT of the last S1 and the AT of S2 (DI=AT_S2_ – RT_S1_). ARI of the S2 beat was calculated as the difference in RT and AT of the S2 (ARI_S2_=RT_S2_ – AT_S2_). **(H)** Representative pre- and post-TEA restitution curves derived from pacing from the RVOT at progressively shorter coupling intervals. Maximum slope of the ARI restitution curve (S_max_) was significantly decreased post-TEA at both the **(I)** RVOT and **(J)** LV border zone. STIM=channel used for stimulation/pacing. Pacing thresholds and electrical restitution (S_max_) were compared pre-TEA *vs* post-TEA by Wilcoxon rank signed test. Pre-TEA *vs* post-TEA ERPs were compared by Student’s paired *t*-test. *N*=10 animals for RV endo and RVOT ERP. *N*=5 for RVOT electrical restitution. *N*=4 for pacing threshold, LV border zone ERP and electrical restitution.

### Effects of TEA on cardiac autonomic function

To confirm that TEA did not affect intrathoracic efferent autonomic function distal to the blockade site, the stellate ganglia and cervical vagi were electrically stimulated pre- and post-TEA (*N=*6), Supplemental Figure 4. There were no significant differences in either the threshold of stimulation or the amplitude of stimulation effects on ventricular ARIs before *vs* after TEA.

We then investigated whether TEA would have any effect on vagal baroreflex sensitivity and vagal tone. Hence, we administered the alpha-adrenoreceptor agonist phenylephrine and assessed the baroreflex response (*N*=10) pre- and post-TEA, Figure 5A-B. Despite a lower phenylephrine-induced blood pressure response post-TEA (44.8±1.1 mmHg pre-TEA *vs* 39.7±1.8 mmHg post-TEA; p=0.034), the heart rate response was significantly greater after TEA (pre-TEA −2.8±1.9 bpm *vs* post-TEA −6.6±2.6 bpm; p=0.0017), Figure 5C-D. Assessing the beat-to-beat relationship of systolic pressure to the RR interval further revealed that TEA significantly increased the baroreflex sensitivity in these chronically infarcted animals from 0.47±0.26 mmHg/msec to 1.41±0.39 mmHg/msec (p=0.0017), Figure 5E.

**Figure 5.**
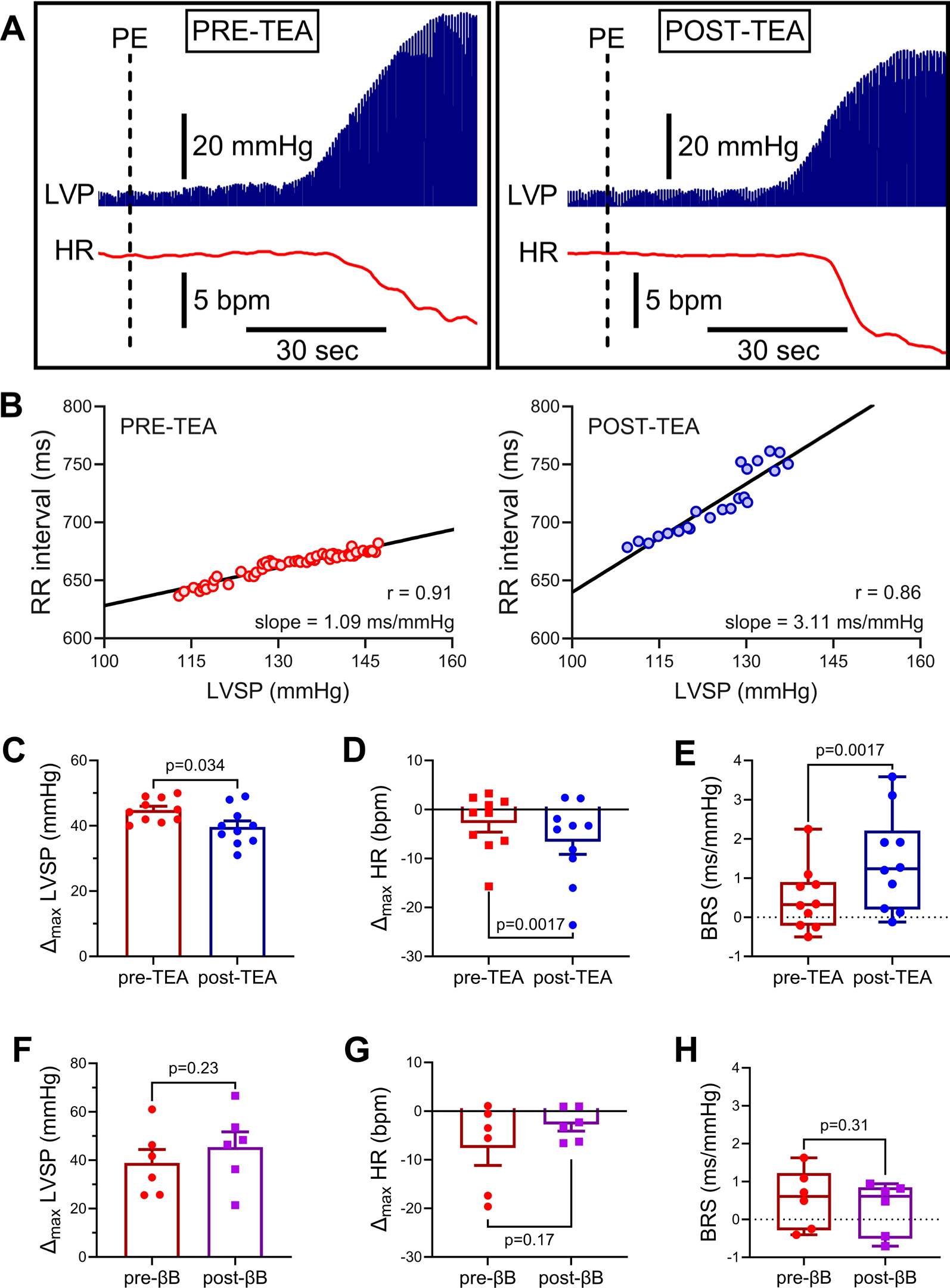
Effects of thoracic epidural anesthesia on baroreflex sensitivity. **(A)** Representative raw trace depicting changes in heart rate (HR) and left ventricular pressure (LVP) upon phenylephrine infusion before and after thoracic epidural anesthesia (TEA). The same dose of phenylephrine was used pre and post TEA. **(B)** Beat-to-beat changes in left ventricular systolic pressure (LVSP) and RR interval were plotted to assess baroreflex sensitivity (BRS) before and after TEA. Although phenylephrine caused **(C)** a slightly lesser increase in peak systolic pressure after TEA, **(D)** a greater slowing of heart rate after TEA was observed. **(E)** BRS, the beat-to-beat relationship between systolic pressure and RR interval, was significantly increased after TEA. Phenylephrine-induced changes in **(F)** LVSP and **(G)** heart rate were similar before and after administration of metoprolol (5 mg). **(H)** Metoprolol did not alter BRS. PE=phenylephrine. Peak change in LVSP and HR pre-intervention *vs* post-intervention (TEA or metoprolol) was compared by paired Student’s *t*-test and BRS was compared by paired Wilcoxon signed ranked test, *N*=10 animals for pre- *vs* post-TEA. *N*=6 for pre- *vs* post-metoprolol.

Given that the effects of TEA are canonically mediated via inhibition of spinal sympathetic outflow,^38^ we sought to determine whether the improvement in BRS was at least partially mediated by reductions in sympathetic efferent tone. To this end, in a separate group of animals (*N*=6), we measured BRS pre and post administration of the β-adrenergic receptor blocker metoprolol (5 mg, IV; 0.096±0.002 mg/kg). After IV metoprolol, however, none of the metrics of phenylephrine-induced baroreflex (absolute change in heart rate or slope of the RR interval to beat-to-beat blood pressure) were improved, Figure 5F-H.

Lastly, to confirm that the effects of TEA may, at least in part, be mediated through augmentation of central vagal drive, we evaluated the activity of parasympathetic neurons within the intrinsic cardiac nervous system by obtaining direct neural recording from the VIVGP (*N*=6, Figure 6A-D). The VIVGP has been shown to predominantly house the somata of post-ganglionic parasympathetic neurons that innervate the cardiac ventricles,^39,40^ overlapping largely with the vascularization of the anterior wall of the ventricles by the LAD coronary artery, which was the region infarcted in this study. These neurons have been previously shown to be silenced or undergo changes in their activity that are consistent with receiving decreased vagal inputs post-MI.^25,39^

**Figure 6.**
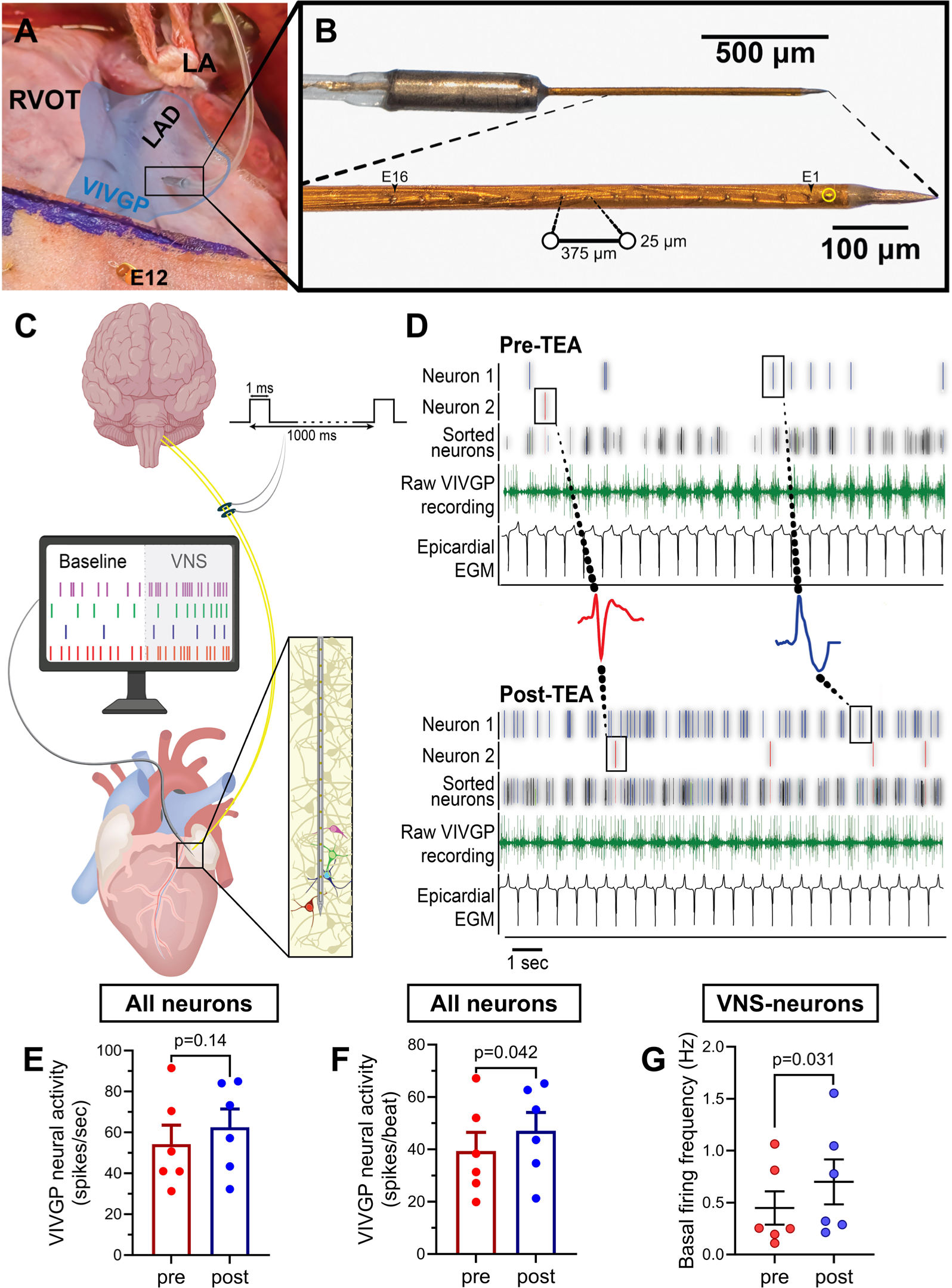
In vivo neural recordings from the ventral interventricular ganglionated plexus (VIVGP). **(A)** Representative image of VIVGP neural recording during terminal experiments. The left atrial appendage (LA) is retracted to expose the VIVGP (shaded blue region), which encircles the base of the left anterior descending coronary artery (LAD). The right ventricular outflow tract (RVOT) is noted for further anatomical reference. The neural recording electrode is inserted at an oblique angle into the fat pad. Seen below is the 56-electrode sock with a basal electrode noted (E12). **(B)** Enlarged view of the customized 16-channel linear microelectrode array probe, which was used to record *in vivo* extracellular action potentials from neurons of the VIVGP. **(C)** Schematic representation depicting the workflow for identification of VIVGP neurons that receive vagal input. The Skellam test was used to identify neurons significantly responding to VNS. **(D)** Representative neuronal recordings from the VIVGP demonstrating two postganglionic parasympathetic neurons increasing their firing frequency post-TEA. **(E-F)** Quantified data showed that while TEA did not affect bulk firing per second it increased bulk firing per ventricular beat. **(G)** Furthermore, TEA increased the basal activity of neurons that specifically respond to VNS (e.g., postganglionic parasympathetic neurons). Pre-TEA *vs* post-TEA parameters were compared using Student’s paired *t*-test. *N*=6 animals, *n*=69 neurons.

TEA had no effect on bulk neural activity (54.3±9.2 spikes/sec pre-TEA to 62.5±8.9 spikes/sec post-TEA; p=0.14), Figure 6E. However, taking into consideration that the majority of neurons of the VIVGP are phase-locked to the cardiac cycle,^41^ we normalized this activity for changes in heart rate to better unmask the effects of TEA on VIVGP neural dynamics. After normalizing for the number of heart beats, we observed an increase in bulk neuronal activity from 39.3±7.1 spikes/beat pre-TEA to 47.1±7.0 spikes/beat post-TEA (p=0.042), Figure 6F.

While total bulk firing within the VIVGP may be suggestive of an enhanced capacity for beat-to-beat neural responses and accommodation, the post-ganglionic neurons of the VIVGP are not simple relay stations. In fact, while the VIVGP houses primarily cholinergic neurons, it also consists of interneurons, and receives an array of inputs from both arms of the autonomic nervous system.^40^ Thus, to confirm whether affected neurons are in fact transducing vagal parasympathetic inputs, we identified unique extracellular neuronal signatures that significantly responded to right and/or left VNS (and hence, likely received vagal inputs), and monitored their activity over time in response to TEA (*N*=6 animals, *n=*69 neurons total). This more granular analysis revealed that the specific activity of these putative efferent post-ganglionic parasympathetic neurons increased by nearly 1.5-fold following TEA (0.45±0.16 spikes/sec pre-TEA to 0.70±0.22 spikes/sec post-TEA; p=0.031), Figure 6G. This along with the BRS data suggested that that TEA improves both afferent and efferent central vagal tone.

## DISCUSSION

### Major findings

In the present study, a comprehensive assessment of the electrophysiological, hemodynamic, and autonomic effects of TEA on cardiac function in a chronic post-infarct model was undertaken to better understand its potential anti-arrhythmic and hemodynamic effects. This study demonstrates that the therapeutic effects of TEA in infarcted hearts may not only be due to sympathetic efferent blockade, but are also likely mediated by relieving sympathetic afferent-mediated suppression of central vagal outflow. Hence, by blocking sympathetic afferent neurotransmission, TEA improves afferent and efferent vagal function, as determined by BRS and VIVGP neuronal recordings, a novel finding of this study. Improved vagal function in addition to sympathetic efferent blockade can lead to increased electrical stability and reduction in the incidence of ventricular arrhythmias by increasing ventricular ERP and ARIs and decreasing restitution slope. Finally, TEA increases atrial ERP and AH interval in infarcted hearts without effects on HV interval, and, notably, did not have significant detrimental effects on RV function, as previously postulated.

### Safety of TEA after myocardial infarction

A major advantage of TEA for the treatment of ventricular arrhythmias is that, unlike intubation and sedation, it allows for the patient to be awake, participating in their medical care, and unlike stellate block, can block afferent and efferent signaling to and from *bilateral* sympathetic chains simultaneously. However, safety data regarding TEA in diseased hearts in both humans and animals are lacking, and a mild decrease in LV contractility was previously reported with TEA in normal ventricles,^42^ resulting in further concerns about the hemodynamic effects of TEA in post-infarct hearts.^43^ In this study of chronically infarcted pigs, a modest reduction in LV inotropy and lusitropy was observed, without overt hemodynamic collapse or compromise.

Moreover, previous studies in healthy hearts and in patients with pulmonary hypertension had suggested a potential decrease in RV preload after TEA, which could potentially compromise RV function or its ability to respond to the increased afterload in pulmonary hypertension.^9,44,45^ Data on the effects of TEA on RV function in the setting of LV dysfunction are lacking. Our careful assessment of RV function parameters in this study demonstrated that TEA did not affect RV systolic pressure or inotropy, and suggests that in the setting of MI, the use of TEA as an anti-arrhythmic therapy is likely to be both effective and safe.

### Electrophysiological effects of TEA

Studies investigating the electrophysiological effects of TEA in infarcted hearts are limited in both scope and quantity. Electrophysiological data from prior studies are largely limited to surface electrocardiogram (ECG) recording in patients or animals with normal hearts and showed mixed results on parameters such as RR and QT interval, with some studies reporting no effect, while others noting shortening of these ECG parameters.^46–48^ These prior studies may be limited by the fact that the benefits of neuromodulation may be masked in healthy animals,^49^ thereby hindering the extrapolation of results from normal animals and humans to those with diseased hearts, where sympathoexcitation and neural remodeling are more prominent. Furthermore, data on the effects of TEA on the conduction system were lacking. This study demonstrated that TEA prolonged the AH interval without altering the HV interval. The observed increase in AH interval may be driven by both blockade of sympathetic efferent neurotransmission as well as improvement in vagal efferent tone. We also found that atrial ERP significantly increased after epidural anesthesia with lidocaine. This data is in accordance with meta-analyses reporting that TEA reduces the incidence of supraventricular arrhythmias after cardiac surgery.^50–52^ Although the goal of this study was not to test the value of this therapy in the setting of atrial fibrillation (AF), the data are hypothesis generating and suggest that further studies looking at the benefit of TEA on AF occurrence in the setting of structural heart disease would have a biological rationale.

This is also the first study to examine the effects of TEA on action potential duration, as measured by detailed electrophysiological mapping in structurally diseased hearts. In line with prolonging ventricular refractoriness, TEA also prolonged ventricular action potential duration with effects observed across viable, scar, and border zone regions of the ventricles. Notably, while TEA prolonged sinus rhythm ARIs by approximately 20 msec, ERPs measured from the same region prolonged by nearly twice this amount, suggesting that the benefits of TEA may be underestimated by changes in the basal state and unmasked by stressors, such as ventricular extra-stimulus pacing. Importantly, TEA mitigated ARI heterogeneity within the border zone regions. Altogether, prolongation of action potential duration and ERP coupled with reductions in electrical heterogeneity in the border zone regions, which are known to house critical sites for reentrant circuits and premature ventricular contractions that trigger ventricular tachycardia/fibrillation,^35–37^ suggests that in chronically diseased hearts, TEA may act at these sites of myocardial and neural heterogeneity^10,53^ to increase refractoriness and suppress arrhythmias.

Furthermore, prior studies in normal canine hearts suggested that effects of TEA on ventricular repolarization in healthy hearts may be greater at faster pacing cycle lengths,^54^ suggesting that the effects observed in this study at resting rates may be an underestimation of the antiarrhythmic potential of TEA. With serially reducing the extra-stimulus coupling interval, we were further able to dissect these dynamics and examine electrical restitution at several sites. A steep restitution slope is reflective of unstable wave propagation that culminates in wave break, ultimately precipitating the initiation of VT/VF.^55^ Importantly, sympathoexcitation steepens electrical restitution and these effects are opposed by parasympathetic activation.^4^ In line with its anti-arrhythmic effects, TEA flattened the slope of the ventricular restitution curves, including in border zone regions.

Combined, the electrophysiological effects of TEA led to an overall reduction in inducibility of ventricular tachyarrhythmias by nearly 70%. This reduction in inducibility of VT/VF is comparable to the antiarrhythmic benefit of TEA reported in case series of patients with VT storm.^6^ This benefit likely results from increased ventricular electrical stability, driven simultaneously by both sympathetic blockade and augmentation of central vagal drive.

### Effects of TEA on parasympathetic function

MI leads to significant parasympathetic dysfunction, which manifests as decreased BRS.^56^ Reduced BRS is an independent predictor of sudden cardiac death and ventricular arrhythmias in patients with MI and heart failure.^56,57^ Previous studies have suggested that the effects of TEA are predominantly due to blockade of sympathetic efferents.^58^ However, our study also suggests that at least a portion of the anti-arrhythmic benefit of TEA may be due to enhanced vagal function.

Studies in healthy hearts had suggested that TEA may not change or may even mildly reduce responses to phenylephrine.^59,60^ Effects of TEA on BRS in humans or animals with diseased hearts remained unknown. As expected, baseline BRS in the chronically infarcted pigs in this study was low. However, TEA led to a significant improvement in BRS in pigs after chronic MI. These data suggest that in chronically infarcted hearts, where sympathetic hyperactivity acts in tandem with parasympathetic dysfunction, TEA significantly improves vagal function.

Correspondingly, increasing sympathetic afferent activity with application of epicardial capsaicin or electrical stimulation of the central end of the left cardiac sympathetic nerve was reported to blunt the baroreflex in rats with chronic heart failure,^61^ and reduced recorded vagal activity in cats with normal hearts.^62^ Notably, in this study, similar beneficial effects on BRS were not observed by blocking sympathetic beta-adrenergic receptors with intravenous metoprolol. This data suggests the beneficial effects of TEA on BRS are less likely to be driven by the decrease in sympathetic efferent tone, and more likely to be due to blockade of spinal afferent neurotransmission. Previous studies in animals have shown MI leads to sympathetic afferent as well as efferent activation.^49^ Hence, by blocking these afferents, TEA may relieve sympathetic afferent mediated suppression of central vagal drive.

Consistent with these findings, by obtaining neuronal recordings from the VIVGP, we noted increases in bulk neural firing in this ganglionated plexus post-TEA. We further corroborated these findings by analyzing the activity of individual neurons receiving vagal inputs, which also increased with TEA. This data provides more direct evidence that TEA improves central efferent vagal drive to post-ganglionic parasympathetic neurons innervating the ventricles. Hence, in the setting of chronic MI where both sympathetic efferent and afferent activation occurs, epidural anesthesia enhances vagal function. Our data further provides important mechanistic support for the sympathetic afferent-mediated reduction of parasympathetic tone following MI (Abstract Illustration). In this regard, it is possible that the anti-arrhythmic, electrophysiological effects of TEA, including increases in ventricular ERP and ARI, are mediated in part through improved central vagal drive. Kock *et al*. demonstrated that TEA may improve LV function in the setting of coronary artery disease, despite maximal beta-blockade,^63^ suggesting an overlooked, non-canonical therapeutic mechanism. Interestingly, the data from our study suggests a potential therapeutic role for targeted sympathetic afferent blockade, which can not only decrease sympathetic efferent tone, but can improve vagal tone. Several studies haves shown that modest levels of vagal activation can reduce ventricular arrhythmias in both diseased and normal porcine and canine hearts.^10,25,64,65^

### Clinical implications of TEA

This study demonstrates that TEA is a safe and effective anti-arrhythmic therapy in a large animal model of chronic MI. Clinically, TEA is a favorable intervention, as it can be instituted at the bedside without specialized equipment, and its effects are rapid. TEA can be used as a continuous infusion as a bridge to more definitive therapies, such as catheter ablation or cardiac sympathetic denervation, allowing for stabilization of patients prior to these more invasive procedures. Furthermore, given that intubation and sedation are first line therapies for the treatment of VT storm refractory to anti-arrhythmic drugs, TEA can allow for weaning of sedation and extubation of the patient, permitting their participation in their own treatment in a shared decision-making process. Nevertheless, the wide use of blood thinners in cardiac patients might pose a limitation, and general contraindications to epidural catheter placement remain (e.g., bacteremia, increased cranial pressure).

### Limitations

General anesthesia with isoflurane is known to blunt autonomic responses. To limit this effect, once surgical procedures were completed, anesthesia was switched to α-chloralose, and data acquisition was initiated after hemodynamic parameters had stabilized for at least one hour. In this study, systemic lidocaine levels were not measured, and it is possible that a minimal amount of lidocaine could have leaked into the systemic circulation, potentially reducing VT inducibility. This, however, is unlikely, as systemic lidocaine has been shown to reduce action potential duration by blocking a portion of the activated sodium channels that does not inactivate during the plateau phase of the action potential,^66,67^ while in this study, prolongation of cardiac action potential durations was observed.

Previous studies have observed similar degrees of neuronal remodeling regardless of infarct site,^26^ suggesting that neuromodulation may be beneficial regardless of the site of infarct. This study, however, demonstrates the efficacy and safety of TEA in the setting of anterior and apical RV and LV infarcts. Whether similar effects on RV hemodynamics would be observed in the setting of posterior/inferior MI was not examined in this study and requires additional investigation.

Finally, the effects of TEA reported in this study may be underestimated by possibility of incomplete cardiac spinal nerve blockade. However, the blockade, herein, was sufficient to achieve a decrease in VT inducibility.

## CONCLUSIONS

This study demonstrates that TEA is effective in reducing ventricular arrhythmias post-MI, without compromising RV function. TEA increases atrial and ventricular ERPs and ARIs, flattens ventricular restitution, and decreases electrical heterogeneity within border zones regions, the likely mechanisms behind TEA’s anti-arrhythmic effects. Notably, while TEA has been long purported to suppress cardiac sympathetic efferent outflow, this study demonstrates that TEA also improves vagal function in diseased hearts, which can independently improve electrophysiological parameters and reduce arrhythmias. The effects of TEA on BRS and parasympathetic neural activity provide novel insights into parasympathetic dysfunction post-MI, demonstrating that blocking sympathetic spinal afferents improves central cardiac vagal drive. TEA uniquely benefits from blockade at an anatomic and autonomic nexus point, achieving bilateral blockade of cardiac-projecting sympathetic efferents along with sympathetic afferents, which may reduce further sympathoexcitation and simultaneously relieve afferent-mediated suppression of parasympathetic outflow.

## NOVELTY & SIGNIFICANCE

### What is known?

- Myocardial infarction induces autonomic dysfunction characterized by sympathetic hyperactivity and parasympathetic withdrawal.
- Thoracic epidural anesthesia has been shown to affect myocardial contractility in normal hearts, raising concerns for safety, but reported to reduce burden of VT in very small case series of patients with refractory arrhythmias.

### What new information does this article contribute?

Thoracic epidural anesthesia suppresses ventricular arrhythmogenesis through blockade of sympathetic efferent as well as spinal sympathetic afferents, which (1) can reduce subsequent reflex sympathetic efferent neurotransmission, and (2) relieve sympathetic inhibition of central vagal tone, improving baroreflex sensitivity and post-ganglionic parasympathetic neural activity. The anti-arrhythmic benefits of TEA are mediated by increases in ventricular effective refractory period and myocardial action potential duration, flattening of ventricular restitution, and mitigation of action potential dispersion in border zone regions. The augmentation of parasympathetic function likely contributes to the observed electrophysiological benefits, potentially pointing to an important additional target for neuromodulation: the blockade of sympathetic afferent neurotransmission to improve vagal tone and reduce subsequent sympathetic efferent tone, and hence, ventricular arrhythmias.

## DATA & SOFTWARE AVAILABILITY

The data for the manuscript is available upon reasonable request from the corresponding author. ScalDyn is freely available upon request from Robert L Lux.

## ACKNOWLEDGMENTS

We would like to thank the UCLA Cardiac Arrhythmia Center faculty for their support.

## AUTHOR CONTRIBUTIONS

JDH, VVW, KK, RLL and MV contributed to conception and design of experiments. JDH, VVW, KK, MAS, CAC, NRJ, ZAL and MV performed experiments and JDH, VVW, RLL and MV analyzed the data. JDH and MV drafted the manuscript and all authors revised and approved the final version of this manuscript.

## Non-standard Abbreviations and Acronyms

AH: atrio-His conduction
ARI: activation recovery interval
AT: activation time (of the EGM)
BRS: baroreflex sensitivity
BSS: bilateral stellate ganglia stimulation
CL: cycle length
DI: diastolic interval
d*P*/d*t*: contractility
ECG: electrocardiogram
EGM: electrogram
ERP: effective refractory period
HV: His-ventricle conduction
ICNS: intrinsic cardiac nervous system
IM: intramuscular
IV: intravenous
LAD: left anterior descending coronary artery
LV: left ventricle
MI: myocardial infarction
MMVT: monomorphic ventricular tachycardia
NSVT: non-sustained ventricular tachycardia
PMVT: polymorphic ventricular tachycardia
RT: recovery time (of the EGM)
RV: right ventricle
RVOT: right ventricular outflow tract
TEA: thoracic epidural anesthesia
VF: ventricular fibrillation
VIVGP: ventral interventricular ganglionated plexus
VNS: vagal nerve stimulation
VT: ventricular tachycardia
VT/VF: ventricular tachyarrhythmias

